# Explore the mechanism of Zigui Yichong Formula in reducing the apoptosis of ovarian granulosa cells in premature ovarian insufficiency based on network pharmacology, molecular docking and cell experiments

**DOI:** 10.1101/2024.09.21.614279

**Authors:** Xin-Miao Zhang, Hong-Yan Xi, Tian-Yu Gao, Shu-Peng Liu, Rong-Xia Li

## Abstract

**Objective:** This research is conducted with the objective of exploring the underlying mechanism by which the Zigui Yichong Formula (ZGYCF) diminishes granulosa cell apoptosis in the context of premature ovarian insufficiency (POI), utilizing network pharmacology, molecular docking, and cellular experimentation approaches.

**Methods:** The active constituents and potential therapeutic targets of the 12 medicinal herbs in ZGYCF, which include Rehmannia glutinosa, Cervus nippon, Cornus officinalis, Ligustrum lucidum, Lycium barbarum, Paeonia lactiflora, Astragalus membranaceus, Codonopsis pilosula, Atractylodes macrocephala, Angelica sinensis, Cyperus rotundus, and Glycyrrhiza uralensis, were identified through searches in the TCMSP, BATMAN, HERB, and ETCM databases. Targets associated with the POI condition were gathered from the OpenTargets, DrugBank, and GeneCards databases. Subsequently, a Venn diagram illustrating the compound-target-disease interaction was generated to derive a set of common targets that bridge the gap between pharmacological and pathological targets. A drug-component-target-disease network diagram was created using Cytoscape 3.9.1. Additionally, protein-protein interaction (PPI) networks were built utilizing the STRING database and visualized with Cytoscape to pinpoint key targets within the overlapping target set. Functional annotation and pathway enrichment analyses, including GO and KEGG pathway analyses, were performed using the clusterProfiler package in R 4.2.1 to investigate the underlying mechanisms by which the drug may influence the disease state. The molecular docking of pivotal active constituents with central targets was carried out using AutoDock Tools. Following this, in vitro studies were executed to corroborate the anticipated mechanisms of action of ZGYCF on POI that were inferred from the network pharmacology analysis.

**Results:** The selected active components include quercetin, kaempferol, and *β*-sitosterol. The core targets identified are Tp53, Bcl-2, and Caspase-3. GO functional and KEGG enrichment analyses indicate that these core targets are primarily enriched in the p53 signaling pathway. Molecular docking results show that quercetin, kaempferol, and *β*-sitosterol have good binding affinity with TP53, Bcl-2, and Caspase-3. Additionally, in vitro experiments demonstrate that ZGYCF medicated serum can reduce ACR-induced apoptosis in KGN cells, increase Bcl-2 expression, and decrease the expression of p53, Bax, Caspase-3, and the Bcl-2/Bax ratio.

**Conclusion:** ZGYCF exerts therapeutic effects on POI through multiple targets and pathways, and it may reduce ACR-induced apoptosis in KGN cells by modulating the p53 signaling pathway.

**Graphical abstract:** 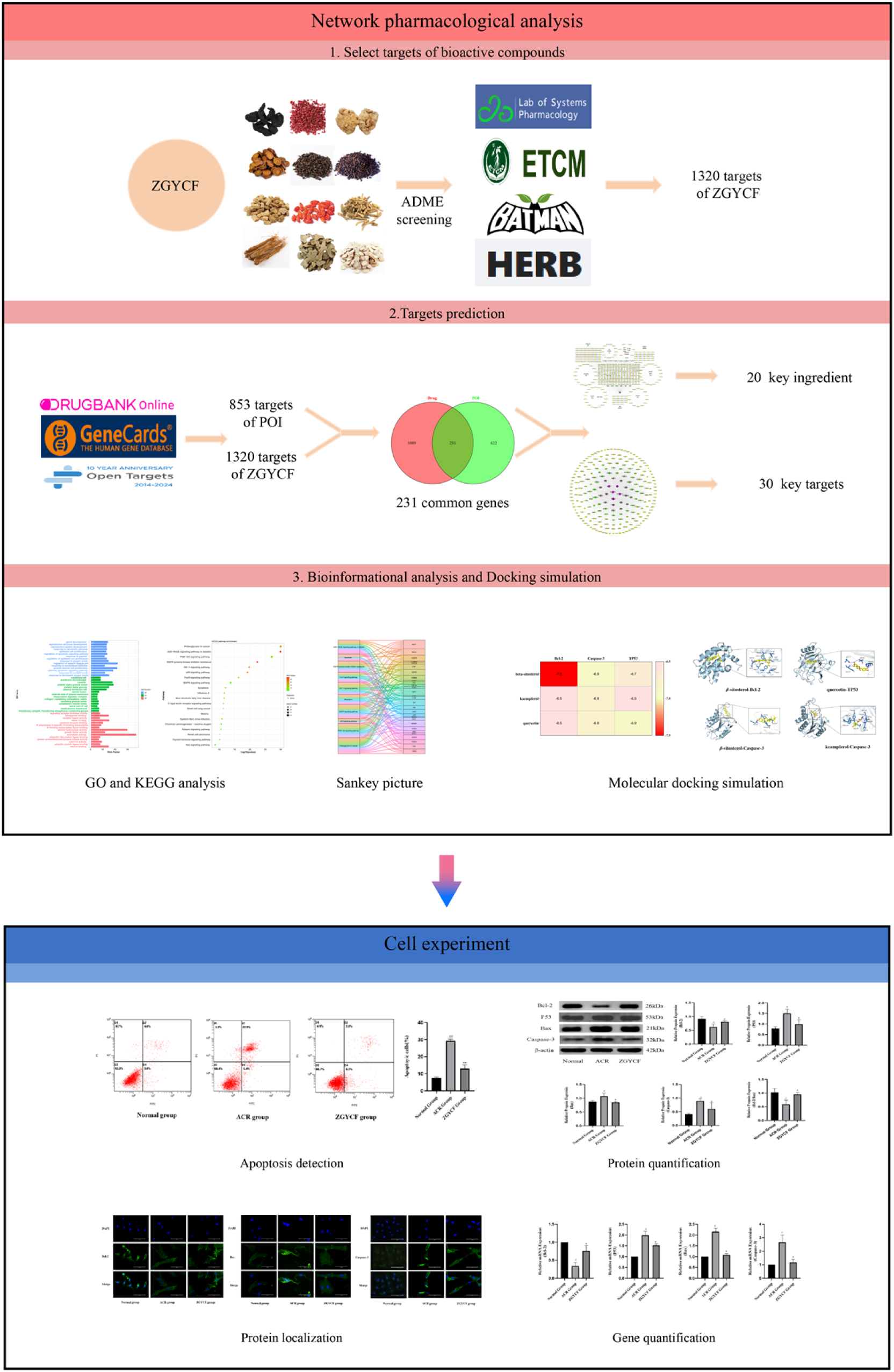

## 1 Introduction

Premature ovarian insufficiency (POI) is a medical condition characterized by the the deterioration of ovarian functionality before reaching 40 years of age. Its main symptoms include erratic or missing menstrual periods, heightened levels of gonadotropins (FSH > 25 mIU/ml on two occasions, with a 4-week interval), and a reduction in estrogen levels^[1]^. Global epidemiological studies indicate that the incidence of premature ovarian insufficiency in females up to 40 years old is roughly 1%, whereas in China, the prevalence is around 2.8% for women in the same age group^[2]^. In recent years, with socio-economic development and changes in living environments, the incidence of POI has been increasing. Certain chemicals in the environment, such as acrolein, can have toxic effects on human reproduction^[3]^. Research indicates that women residing in low-and middle-income nations are commonly subjected to household cooking fumes and passive smoke, resulting in significant inhalation of acrolein. Such exposure may induce transgenerational ovarian impairment and hasten the aging process of the ovaries, which could precipitate or exacerbate the development of POI^[4]^. However, there are currently no effective treatments for POI. Therefore, it is crucial to identify safer and more effective alternative treatment options.

Traditional Chinese medicine (TCM) offers unique advantages in treating POI. Zigui Yichong Formula (ZGYCF), a clinical prescription from professor Du Huilan, a guiding teacher of the sixth and seventh batches of the national TCM experts’ academic experience inheritance program, consists of Rehmannia glutinosa, Cervus nippon, Cornus officinalis, Ligustrum lucidum, Lycium barbarum, Paeonia lactiflora, Astragalus membranaceus, Codonopsis pilosula, Atractylodes macrocephala, Angelica sinensis, Cyperus rotundus, and Glycyrrhiza uralensis. It is effective in nourishing the kidneys and filling essence, soothing the liver, and regulating the spleen, showing significant clinical efficacy in treating ovarian dysfunction^[5]^. Preliminary animal studies have confirmed that ZGYCF can effectively improve serum hormone levels in rats with reduced ovarian reserve, reduce follicle atresia, and protect ovarian reserve function^[6]^. However, its biological components, pharmacological effects, and molecular mechanisms have not yet been fully elucidated.

Network pharmacology, a burgeoning discipline, draws upon principles from systems biology, genomics, proteomics, and related fields. It employs high-throughput omics data analysis, computational modeling, and network database queries to uncover the intricate network connections among drugs, chemical constituents, biological targets, and diseases. By analyzing these network relationships, network pharmacology predicts drug mechanisms of action, evaluates drug efficacy and adverse effects, and aims to identify highly effective and low-toxicity drugs^[7]^. Molecular docking is a form of computational modeling that simulates the interactions between molecules and proteins on an atomic scale. It forecasts the possible shapes that ligands and receptors might adopt and computes various parameters, including binding affinity, to evaluate the strength of their interactions. This technique helps reveal the pharmacological mechanisms of herbs and aids in predicting the efficacy, quality, and formulation of TCMs^[8, 9]^. To this end, we use a combination of network pharmacology, molecular docking, and experimental methods to explore the effective components, potential targets, and molecular mechanisms of ZGYCF in treating POI, providing a reference for discovering new treatment options for POI.

## 2 Materials

### 2.1 Experimental cells and animals

The human ovarian granulosa cell line (KGN, CL-0603) was purchased from Procell Life Technology Co., Ltd. (Wuhan, China). Thirty SPF-grade healthy female SD rats, weighing (200±20) g, were obtained from Vital River Laboratory Animal Technology Co., Ltd. (Beijing, China), with an animal production license number SCXK (Jing) 2021-0006. The rats were housed at the SPF Laboratory Animal Center of Hebei University of Chinese Medicine, with a temperature of (24±2) ℃, humidity of 40% to 70%, and a light cycle from 08:00 to 20:00, with free access to water and food. This experiment was reviewed and approved by the Animal Ethics Committee of Hebei University of Chinese Medicine ( DWLL202302006).

### 2.2 Experimental drugs and reagents

Composition of ZGYCF: Rehmannia glutinosa 20 g (21102341), Placenta hominis 6 g (21072661), Cornus officinalis 12 g (21092031), Ligustrum lucidum12 g (21080771), Lycium barbarum 12 g (21081731), Paeonia lactiflora 10 g (21093011), Astragalus membranaceus 15 g (21081731), Codonopsis pilosula 12 g (21102831), Atractylodes macrocephala10 g (21101341), Angelica sinensis 10 g ( 21092821), Cyperus rotundus 10 g (21061081), Glycyrrhiza uralensis 6 g (21101822). The above medicines are provided as ready-to-use herbal granules by Shenwei Pharmaceutical Co., Ltd. ACR (A12346, purity > 98 %) was procured from Aladdin Biochemical Technology Co., Ltd. (Shanghai, China). B-cell lymphoma-2 antibody (Bcl-2, 26593-1-AP), B-cell lymphoma-2 associated X protein antibody (Bax, 60267-1-Ig), and cysteinyl aspartate specific puroteinase-3 antibody (Caspase-3, 66470-2-Ig) were purchased from Proteintech (Wuhan, China), and the p53 antibody (Ab131442), goat anti-rabbit fluorescent secondary antibody (SA00013-2), and goat anti-mouse fluorescent secondary antibody (SA00007-3) were purchased from Abcam (Cambridge, UK). The annexin V-fluorescein isothiocyanate (FITC)/propidium iodide (PI) apoptosis detection kit (20230301) was purchased from CytoScen Innovation Biotech Co., Ltd. (Beijing, China). The RNA extraction kit (R6831-01) was purchased from Omega Bio-Tek ( Georgia, USA). MonScriptTMRTⅢALL-in-OneMix with dsDNase (MR05101M) and MonScriptTMChemoHS qPCR Mix (MQ00401S) were purchased from Mona Biotechnology Co., Ltd. (Suzhou, China). The primers were designed and synthesized by Servicebio Technology Co., Ltd. (Wuhan, China).

## 3 Methods

### 3.1 Screening of active components and target genes of ZGYCF

The constituents of ZGYCF, comprising Rehmannia glutinosa, Placenta Hominis, Cornus officinalis, Ligustrum lucidum, Lycium barbarum, Paeonia lactiflora, Astragalus membranaceus, Codonopsis pilosula, Atractylodes macrocephala, Angelica sinensis, Cyperus rotundus, and Glycyrrhiza uralensis, were identified through searches conducted in the TCMSP (https://old.tcmsp-e.com/tcmsp.php), BATMAN (http://bionet.ncpsb.org.cn/batman-tcm/index.php), HERB (http://herb.ac.cn/), and ETCM (http://bionet.ncpsb.org.cn/batman-tcm/index.php) databases. These resources facilitated the elucidation of the herbal formula’s pharmacologically active ingredients. Subsequent to the initial identification, the components were refined by applying filters for Oral Bioavailability (OB) of at least 30% and Drug Likeness (DL) of no less than 0.18. This process was undertaken to pinpoint the bioactive constituents of ZGYCF along with their associated target data. The UniProt database was employed to harmonize the nomenclature of drug targets and eliminate any redundant entries, resulting in the finalization of the active components and their respective targets.

### 3.2 Screening of POI disease-related targets

The OpenTargets (https://www.opentargets.org/), DrugBank (https://go.drugbank.com/), and GeneCards (https://www.genecards.org/) databases were used to search for disease-related targets. The keyword “premature ovarian insufficiency, POI” was used to identify disease-related targets. The gene names of the disease targets were standardized and duplicates were removed, ultimately obtaining the potential targets for the disease. The disease-associated targets and the pharmacological targets of ZGYCF were uploaded into the Venny 2.1 tool to determine the overlapping targets, subsequently generating a Venn diagram to visualize these intersections.

### 3.3 Construction and analysis of the drug-compound-target-disease network

The interactions between the active constituents and their respective targets of ZGYCF were inputted into Cytoscape 3.9.1 to build a drug-compound-target-disease network diagram. In this network, nodes signify the compounds and targets, with edges depicting the connections and interactions between them. The Network Analyzer plugin was utilized to examine the network’s structure and to compute the degree centrality of the nodes.

### 3.4 Construction of the protein-protein interaction network and screening of core targets

The common targets of ZGYCF’s core drugs and POI were uploaded to the STRING database, with the species restricted to “Homo sapiens”, the minimum required interaction score was set to ≥0.7 (highest confidence), while other parameters remained unchanged. A PPI network was obtained, and the files were saved in both TSV and PNG formats. The obtained PPI information was imported into Cytoscape software in TSV format for visualization, and isolated targets without interactions were removed. The network topology was analyzed, and core targets were selected based on degree ranking (CC/BC/degree ranking) according to the topological structure. The MCC algorithm in the cytoHubba plugin was also used to screen for core targets.

### 3.5 GO function and KEGG pathway enrichment analysis

The pivotal targets associated with the principal constituents of ZGYCF and the pathophysiology of the disease underwent gene ontology (GO) functional enrichment and Kyoto Encyclopedia of Genes and Genomes (KEGG) pathway enrichment analyses via the R clusterProfiler version 4.6.2. In the GO enrichment analysis, a significance level of P < 0.05 was applied as the threshold for selection. The top 20 most significant biological processes with P < 0.05 were then chosen for representation in a bar graph. Similarly, for KEGG enrichment analysis, P < 0.05 was set as the threshold for significant enrichment, and the top 20 pathways with P < 0.05 were selected to generate a bubble chart for visualization.

### 3.6 Molecular docking

Perform molecular docking of the chosen principal compounds against the central targets. The 3D structures of the primary active compounds within the medication were obtained from the PubChem database, while the 3D structures of the key target proteins were sourced from the Protein Data Bank (https://www.rcsb.org/). The central target proteins were first refined using PyMOL 2.6.0 to eliminate solvent molecules. Subsequently, they were subjected to additional processing in AutoDock Tools 1.5.7 for the addition of hydrogen atoms and the assignment of partial charges. The processed target proteins, along with the active compounds, were converted into “pdbqt” formatted files. Grid parameters, including position and size, were then defined. The molecular docking procedure was executed with AutoDock Vina, and the outcomes of the docking were rendered for visualization using PyMOL.

### 3.7 Cell experiment

#### 3.7.1 The preparation of medicated serum containing ZGYCF

Thirty healthy female SD rats, aged between 6 to 7 weeks, were selected and acclimatized for 7 days. Following this acclimatization phase, the rats were randomly assigned to two experimental groups: one receiving ZGYCF medicated serum and the other serving as a blank serum control, with 15 animals in each group. The rats in the ZGYCF group were administered ZGYCF via gavage at a dose of 14.175 g/kg^[6]^, while those in the blank group received an equal volume of distilled water. Gavage was conducted twice daily, at 08:00 in the morning and 20:00 in the evening, for 4 consecutive days. Following the last dose, the rats were anesthetized one hour later with an intraperitoneal injection of 2% sodium pentobarbital at a dose of 10 ml/kg. Subsequently, blood samples were drawn from the femoral artery for analysis. Following collection, the blood samples were incubated at room temperature for a period of 2 hours to facilitate clotting. Subsequently, centrifugation was performed at a rate of 3000 r/min for a total of 10 minutes. The supernatant was collected, subjected to a heat inactivation process at 56°C for 30 minutes to neutralize complement proteins. Sterilization was achieved by passing the serum through a 0.22 μm filter. The processed serum was then aseptically sealed and stored at -80°C until required for subsequent experimental procedures.

#### 3.7.2 Cell grouping and administration

KGN cells from passages 3 to 10 were selected and divided into three groups: the control group, the ACR group, and the ZGYCF group. The control group was incubated with 10% blank serum + DMEM/F12 medium (containing 1% penicillin/streptomycin) for 24 hours; the ACR group was incubated with 50 μmol/L ACR + 10% blank serum + DMEM/F12 medium (containing 1% penicillin/streptomycin) for 24 hours; the ZGYCF group was incubated with 50 μmol/L ACR + 10% ZGYCF medicated serum + DMEM/F12 medium (containing 1% penicillin/streptomycin) for 24 hours.

#### 3.7.3 Annexin V-FITC/PI staining was used to detect the apoptosis rate of cells in each group

KGN cells from the respective groups were harvested utilizing trypsin devoid of EDTA, subsequently rinsed twice with PBS, and enumerated. An aliquot of approximately 5×10⁴ cells was then resuspended and transferred into a 1.5 mL EP tube, followed by centrifugation at a speed of 2000 r/min for a duration of 5 minutes. The supernatant was decanted, and the cell pellet was resuscitated in 500 μL of 1×Annexin V Binding Buffer. Following resuspension, 5 μL of Annexin V-FITC conjugate and propidium iodide stain were introduced into the mixture, which was then gently vortexed to ensure uniform distribution of the dyes. The cells were incubated at room temperature, protected from light, for 5-15 minutes, and cell apoptosis changes were measured using the FC500MCL flow cytometer (Beckman Coulter, USA).

#### 3.7.4 Immunofluorescence staining was used to detect protein expression in KGN cells from each group

Coverslips with cells seeded in 12-well plates, which had been preserved at -80°C, were taken out and permitted to thaw at ambient temperature for a duration of 30 minutes. The coverslips were then subjected to a triple washing procedure with PBS, with each wash cycle lasting for 5 minutes. The coverslips, now bearing the cells, were transferred onto glass slides and delineated with a hydrophobic barrier using a pen. Subsequently, 10% goat serum solution was applied, and the slides were incubated in a water bath at 37°C for 30 minutes to block any nonspecific binding sites. The solution was decanted, and the primary antibody was introduced to the slides, which were then incubated overnight in a moisture-controlled environment at 4°C. Following the overnight incubation, the slides were allowed to equilibrate to room temperature for 30 minutes and subsequently underwent three washes with PBS, each lasting 5 minutes. Fluorescently labeled secondary antibodies were introduced, and the slides were subjected to incubation within a humidified chamber at a temperature of 37°C for a period of one hour, while being safeguarded from exposure to light. After incubation, the slides were washed three times with PBS, each wash lasting 10 minutes, protected from light. The nuclei were stained with DAPI for 5 minutes and then washed five times with PBS, each wash lasting 10 minutes, in the dark. The coverslips were mounted and observed under the AMAFD1000 live-cell imaging system (Thermo Fisher Scientific, USA), protected from light, and photographs were taken.

#### 3.7.5 Western blotting was used to detect protein expression in KGN cells from each group

KGN cells from each group were collected, and 150 µL of protein lysis buffer prepared with RIPA and PMSF was added. The samples were vortexed for 30 minutes and then centrifuged at 8000 r for 10 minutes. The supernatant was collected to obtain the extracted proteins, and the protein concentration was determined using a BCA protein quantification kit. Protein loading buffer was added at a ratio of 4:1. SDS-PAGE gel electrophoresis was performed, followed by transfer onto a PVDF membrane. The membrane was blocked with non-fat milk for 2 hours, then incubated with primary antibodies Bcl-2, p53, Bax, Caspase-3, and *β*-actin (dilutions of 1:2000, 1:2000, 1:5000, 1:2000, and 1:4000, respectively) overnight at 4°C. The corresponding secondary antibodies were added, and the membrane was incubated on a shaker at room temperature for 1 hour. After thorough washing with TBST, imaging was performed using the Touch Imager electronic imaging system (e-BLOT Life Science, Shanghai, China). *β*-actin was used as an internal control, and ImageJ software was used to analyze and generate grayscale values. The relative amount of protein was expressed as the ratio of the optical density of the target bands to the internal control.

#### 3.7.6 Quantitative Real-Time PCR ( qRT-PCR ) was used to detect mRNA expression in KGN cells from each group

KGN cells from each group were collected, and total RNA was extracted using the MicroElute Total RNA Kit according to the manufacturer’s instructions. The RNA purity and concentration were measured. cDNA was synthesized via reverse transcription, and the qPCR reaction system (20 μL) was prepared as follows: 10 μL of qPCR Mix, 0.4 μL of forward and reverse primers each, 0.2 μL of Low ROX Dye, 1 μL of DNA template, and 8 μL of nuclease-free water. Quantification was performed using an ultramicro nucleic acid protein analyzer (Thermo Fisher Scientific, MA, USA). The RNA was pre-denatured at 95°C for 10 minutes, followed by 40 cycles of 95°C for 10 seconds, 60°C for 30 seconds. *β*-actin was used as the reference gene, and the relative mRNA expression levels were calculated using the 2^-ΔΔCt^ method (normalized to the control group). The primer sequences and product sizes are shown in Table 1.

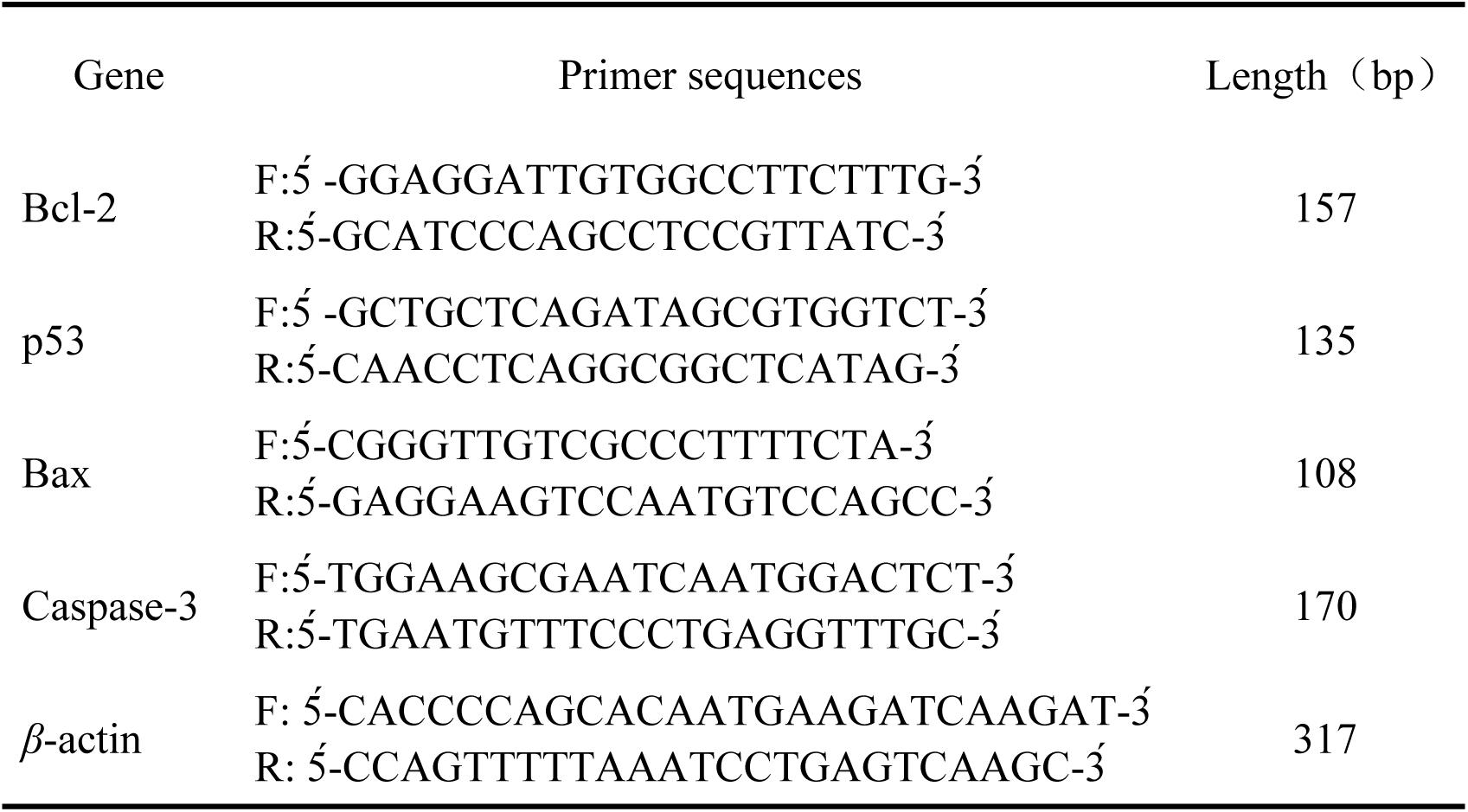

### 3.8 Statistical analysis

Statistical analysis was performed using IBM SPSS 26.0 software. The results were expressed as mean ± SD (x̅±s). For comparisons of multiple groups, ANOVA was used if the variance was homogeneous. Pairwise comparisons between groups were tested using the LSD method. *P* values <0.05 were considered to be statistically significant.

## 4 Results

### 4.1 Active compounds and corresponding drug targets of the core herbs in ZGYCF

The core ingredients of ZGYCF were retrieved from the TCMSP, BATMAN, HERB, and ETCM databases. Active compounds that met the criteria of OB ≥ 30% and DL ≥ 0.18 were screened, resulting in 4 compounds from Rehmannia glutinosa, 2 from Human Placenta, 29 from Cornus officinalis, 19 from Ligustrum lucidum, 73 from Lycium barbarum, 25 from Paeonia lactiflora, 27 from Astragalus membranaceus, 38 from Codonopsis pilosula, 14 from Atractylodes macrocephala, 11 from Angelica sinensis, 20 from Cyperus rotundus, and 3 from Glycyrrhiza uralensis. After removing duplicate components, a total of 211 active compounds were identified. A single herb may contain multiple active compounds, and the same active compound can be present in multiple herbs, indicating that different TCMs share common active compounds. This aligns with the multi-component, multi-target synergistic action mechanism of TCM, where combination usage has a synergistic effect. Potential targets of the active compounds were identified through databases, and after removing duplicates, a total of 1,320 drug targets were obtained (Figure 2a、b).

**Figure 2.**
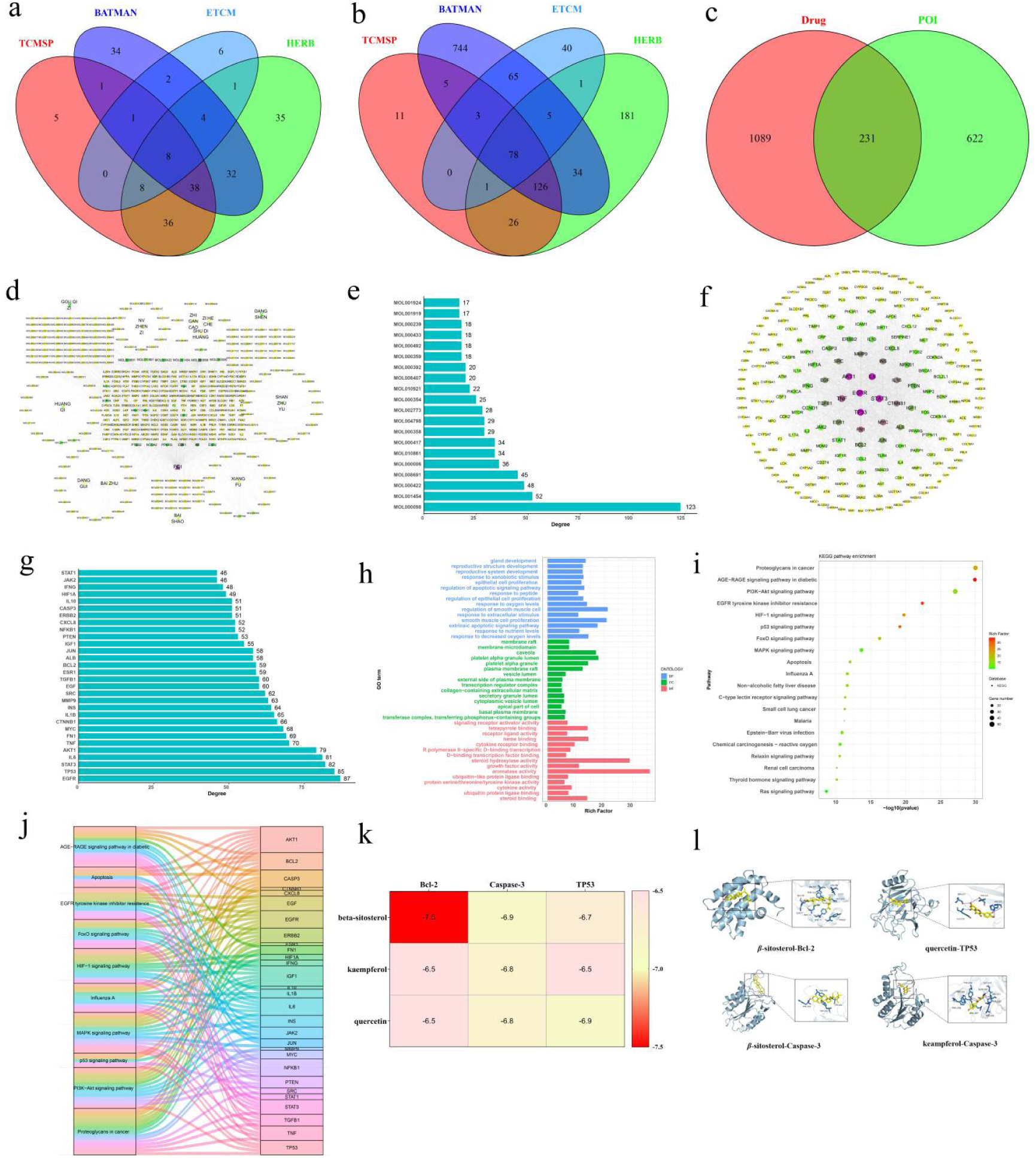
Results of network pharmacology and molecular docking. (a) Intersection of active ingredients of the ZGYCF across different databases. (b) Intersection of targets corresponding to the active ingredients of the ZGYCF across different databases. (c) Venn diagram showing the overlap between drug-targets of the Ziwei Yichong Decoction and disease targets. (d) Network diagram of drug-compound-target-POI for the ZGYCF. (e) Bar chart of the top 20 compounds ranked by connectivity. (f) PPI network diagram. (g) Bar chart of the top 30 core targets ranked by connectivity. (h) GO functional molecular entries chart. (i) Bubble chart of KEGG enrichment analysis. (j) Sankey diagram of the top targets and top pathways. (k) Heatmap of molecular docking binding energies. (l) Molecular docking results between active components and key targets.

### 4.2 The core active ingredient of ZGYCF targets potential pathways for the treatment of POI

The search for POI-related targets was conducted through the OpenTargets database (Score ≥ 0.1), DrugBank (Score ≥ 15), and GeneCards (Score ≥ 15). The results were filtered based on the threshold values, and 853 disease targets were compiled after standardizing and deduplicating the gene names. By intersecting the potential disease targets with the core active ingredient targets of ZGYCF, 231 common targets were identified as potential drug targets for the treatment of the disease (Figure 2c).

### 4.3 Construction and analysis of the Drug-Compound-Target-Disease cetwork

Using the STRING database and Cytoscape, a drug-compound-target-disease network (Figure 2d) was constructed. This network contains 440 nodes and 2,183 edges, where the edges represent the targeting relationships between compounds and targets. Among them, 196 active ingredients act on 231 targets, which to some extent reflects the mechanism of TCM, where different active ingredients can act on the same target, and the same active ingredient can act on multiple targets. The Network Analyzer plugin was used to analyze the topological properties of the network. In the network graph, the size of each node represents its degree value, with a higher degree indicating that the node plays a more central role in the network. The top 20 key components are ranked based on their degree values according to three metrics: degree, betweenness, and closeness (Figure 2e). Based on this analysis, the main active components identified are quercetin, berberine, kaempferol, alpha-carotene/beta,epsilon-carotene, luteolin, Delta-D, calycosin, beta-sitosterol, delphinidin, beta-carotene, and isorhamnetin. These are key nodes in the network, indicating that their anti-POI effects are more significant.

### 4.4 Construction of the PPI network and core target screening

The 231 intersecting drug-disease targets were imported into the STRING database to obtain protein-protein interaction (PPI) relationships, and then imported into Cytoscape software for analysis. The resulting network consists of 223 nodes and 2,411 edges, where the edges represent the interactions between targets, with isolated targets that have no intersections being removed (Figure 2f). Using the Network Analyzer plugin to analyze the network’s topological properties, degree values were calculated, and the color intensity and node size were set to reflect the degree value. A combined screening was performed based on three metrics: degree, betweenness, and closeness, ranking the top 30 core nodes (Figure 2g). The core targets identified are EGFR, Tp53, STAT3, IL6, AKT1, TNF, FN1, MYC, CTNNB1, IL1B, INS, MMP9, SRC, EGF, TGFB1, ESR1, BCL2, ALB, JUN, IGF1, PTEN, NFKB1, CXCL8, ERBB2, and CASP3. These serve as the key targets for the treatment of diseases by the core active ingredients of ZGYCF.

### 4.5 GO function and KEGG pathway enrichment analysis

Based on R clusterProfiler, GO function analysis was performed on the 231 overlapping drug-related disease targets. Bar plots were generated for the top 15 entries in biological process (BP), molecular function (MF), and cellular component (CC) categories, each with a *P*-value < 0.05 (Figure 2h). The vertical axis represents the GO entries, while the horizontal axis indicates the number of targets enriched in each entry. BP is mainly involved in the regulation of apoptotic processes, regulation of epithelial cell proliferation, and regulation of oxygen levels, among others. MF primarily includes enzyme binding, receptor binding, and protease binding, while CC includes cellular organelles such as cytoplasmic vesicles, the external side of the plasma membrane, and membrane rafts. KEGG pathway enrichment analysis was also conducted on the targets, identifying 178 enriched signaling pathways, of which the top 20 pathways with *P*-value < 0.05 were selected for visualization through a bubble chart (Figure 2i). This indicates that ZGYCF prevents POI through various signaling pathways. Combined with the PPI results, Tp53, Bcl-2, and Caspase-3 were found to be enriched in the p53 signaling pathway, suggesting that ZGYCF may improve POI by regulating p53 (Figure 2j).

### 4.6 Molecular docking of the active components of ZGYCF with the core targets of the p53 pathway

Based on the ranking of degree values of active components in the active component-target network and literature review, this study performed molecular docking validation of the active components quercetin, kaempferol, and *β*-sitosterol with the core targets of the p53 pathway, namely Tp53, Bcl-2, and Caspase-3 in the PPI network. The lower the binding energy, the stronger the binding affinity between the core active components and core targets. It is generally considered that core active components with binding energy ≤-5.0 kcal/mol bind well with protein molecules. A heat map was generated based on binding energy (Figure 2k), where deeper colors represent lower binding energies and stronger binding affinities. Conformations with binding energy ≤ -5.0 kcal/mol were visualized (Figure 2l), and the results show that there is a stable interaction between the active components of ZGYCF and the core targets Tp53, Bcl-2, and Caspase-3 with strong binding affinity.

### 4.7 Comparison of apoptosis rates among different groups of KGN cells

In contrast to the normal group, the ACR group exhibited a significant elevation in KGN cell apoptosis (*P*<0.01). Conversely, the ZGYCF treatment group showed a marked reduction in KGN cell apoptosis relative to the ACR group (*P*<0.01) ( Figure 3a、b).

**Figure 3.**
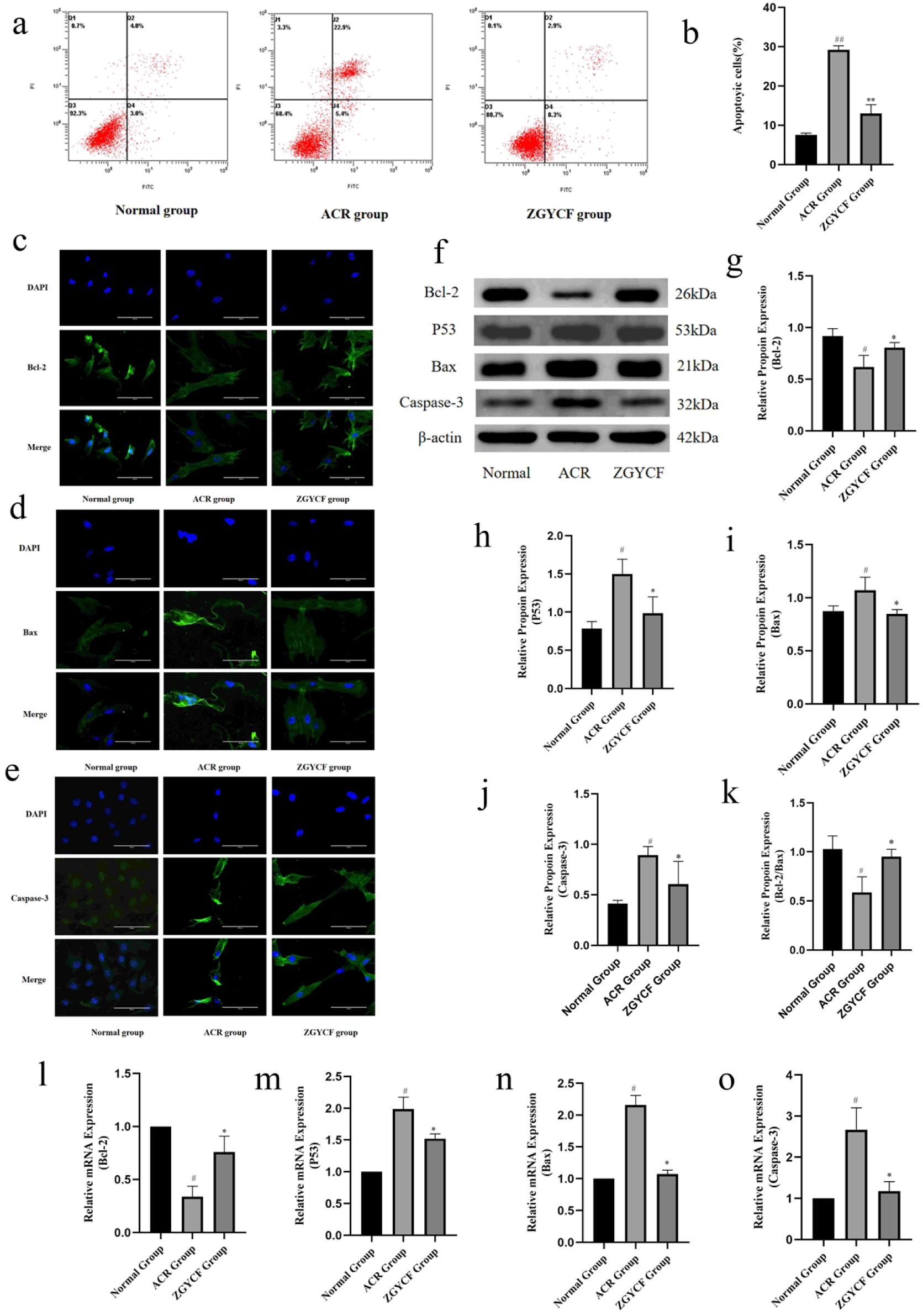
Results of cell experiments. (a) Flow cytometry results of KGN cells from each group. (b) Bar graph comparing apoptosis levels of KGN cells among different groups. (c-e) Immunofluorescence images showing the localization of Bcl-2, Bax, and Caspase-3 proteins in KGN cells from each group (scale bar, 100μm). (f) Western blotting results showing the expression levels of Bcl-2, p53, Bax, and Caspase-3 proteins in KGN cells from each group. (g-k) Bar graphs comparing the relative expression levels of Bcl-2, p53, Bax, and Caspase-3 proteins in KGN cells from each group, as well as the Bcl-2/Bax ratio. (l-o) Bar graphs comparing the relative mRNA expression levels of Bcl-2, p53, Bax, and Caspase-3 in KGN cells from each group. Data are presented as mean ± SD (n = 3). ^#^*P* < 0.05, ^##^*P* < 0.01 compared with the normal group, \**P* < 0.05, \*\**P* < 0.01 compared with the ACR group.

### 4.8 Comparison of Bcl-2, p53, Bax, and Caspase-3 protein expression in KGN cells among different groups

Immunofluorescence was used to detect the expression of apoptotic factors in each group of mice, and it was found that Bcl-2, Bax, and Caspase-3 were predominantly localized in the cytoplasm of granulosa cells (Figure 3c∼e).

In contrast to the normal group, the ACR group demonstrated a significant reduction in the expression of Bcl-2 protein within KGN cells (*P*<0.05), Concurrently, there was an upregulation in the expression of p53, Bax, and Caspase-3 proteins, as well as an increased Bcl-2/Bax ratio (*P*<0.05). In contrast to the ACR group, the expression of Bcl-2 protein was increased in the ZGYCF group (*P*<0.05), while the expression of p53, Bax, and Caspase-3, as well as the Bcl-2/Bax ratio were decreased (*P*<0.05) (Figure 3f∼k).

### 4.9 Comparison of Bcl-2, p53, Bax, and Caspase-3 mRNA expression in KGN cells among different groups

In contrast to the normal group, the mRNA expression of Bcl-2 in KGN cells was observed to decrease in the ACR group (*P*<0.05). Conversely, the mRNA expression of p53, Bax, and Caspase-3 was found to increase (*P*<0.05). In the ZGYCF group, the mRNA expression of Bcl-2 was elevated compared to the ACR group (P<0.05), whereas the expression of p53, Bax, and Caspase-3 was reduced (P<0.05) ( Figure 3l∼o).

## 5. Discussion

During follicular development, granulosa cells provide nutrients and facilitate information exchange for the development and maturation of oocytes through close contact and gap junctions with oocytes^[10]^. Follicular atresia is the main form of follicle and oocyte loss in the ovary, playing a crucial role in maintaining and regulating the homeostasis of the follicular environment and ensuring the development of dominant follicles. However, excessive follicular atresia will accelerate the activation of primordial follicles and reduces ovarian function. The interaction between granulosa cells and theca cells provides estrogen, which can inhibit the expression of pro-apoptotic genes and improve cell survival within the follicle, thus preventing follicular atresia ^[11]^. Tilly et al. first proposed that the essence of follicular atresia is the apoptosis of granulosa cells ^[12]^. Studies have shown that abnormal apoptosis of granulosa cells leads to inappropriate follicular atresia, and when the number of apoptotic granulosa cells in a follicle reaches 10% of the total number of granulosa cells, the follicle enters atresia, thereby triggering the development of POI ^[13, 14]^. With further research into its etiology, it is now believed that abnormal apoptosis of granulosa cells is one of the key causes of POI ^[15]^.

ACR, an α, *β*-unsaturated aldehyde, is prevalent as an environmental contaminant and is frequently detected in foodstuffs. It is introduced into the human system through various routes, including dietary intake, alcohol consumption, smoking, and exposure to fuel combustion emissions^[16]^. Studies have shown that women frequently exposed to household cooking emissions and secondhand smoke can inhale high levels of ACR, leading to ovarian dysfunction and accelerated ovarian aging, which may induce the development of POI ^[4]^. Other studies have indicated that ACR has high cytotoxicity, causing cell activation, detachment, and death ^[17, 18]^. Averill-Bates et al. demonstrated ^[19]^ that ACR can induce apoptosis in Chinese hamster ovarian cells through the intrinsic pathway involving cytochrome c release. In our previous research, we applied ACR at concentrations of 0, 12.5, 25, 50, and 100 μM to KGN cells and found that KGN cell apoptosis increased in a dose-dependent manner^[20]^. Therefore, ACR-induced apoptosis of ovarian granulosa cells may be an important cause of the development of POI.

In TCM literature, POI falls under categories such as “scanty menstruation,” “early cessation of menstruation,” and “infertility.” In 《Su Wen · Yin Yang Ying Xiang Da Lun》 it is recorded: “If one understands the seven damages and eight benefits, then both can be regulated; if one does not use this, early decline will follow. At the age of forty, the yin energy is already half depleted, and physical vitality weakens.” This text was one of the earliest to propose symptoms of early decline and the seven damages, aligning with modern medical understanding of POI. Scholars have various interpretations regarding the etiology and pathogenesis of POI, but they unanimously agree that kidney essence deficiency is the primary pathogenic mechanism of this disease. Additionally, liver stagnation and spleen deficiency are closely related to its occurrence. ZGYCF, a clinical prescription based on professor Du Huilan’s experience, has shown remarkable clinical efficacy in the treatment of ovarian insufficiency-related diseases^[5]^. In this formula, Rehmannia glutinosa and Cervus nippon are the sovereign herbs, replenishing the kidneys, benefiting essence, and nourishing the blood. Cornus officinalis, Ligustrum lucidum, Lycium barbarum, and Paeonia lactiflora serve as minister herbs, tonifying the liver and kidneys, and enriching essence and blood. Astragalus membranaceus, Codonopsis pilosula and Atractylodes macrocephala strengthen the spleen and tonify Qi to assist in blood production, with these six herbs collectively acting as minister herbs. Angelica sinensis and Cyperus rotundus soothe the liver, promote Qi circulation, and nourish and harmonize the blood, ensuring that the tonics do not lead to stagnation, thus serving as assistant herbs. Glycyrrhiza uralensis moderates and harmonizes the spleen and stomach and coordinates the actions of the other herbs, functioning as the envoy herb. When used together, these herbs tonify the kidneys, replenish essence, soothe the liver, and regulate the spleen, harmonizing qi and blood, promoting the smooth flow of the Ren and Chong channels, and ensuring timely menstrual cycles. Our previous research found that ZGYCF can improve serum hormone levels, reduce follicular atresia, and protect ovarian reserve function in POI model mice by regulating mitochondrial energy metabolism and iron metabolism ^[21, 22]^. However, its bioactive components, pharmacological effects, and molecular mechanisms require further investigation. To this end, we used network pharmacology and molecular docking techniques to analyze the effective components, potential targets, and molecular mechanisms of ZGYCF in the treatment of POI.

The combination of network analysis and network database retrieval transforms the traditional “drug-target-disease” model into a new drug development approach, which is suitable for studying the mechanisms of TCM. In this study, through network pharmacology, data mining of the active components and targets of ZGYCF was conducted, resulting in the identification of 1,320 core drug targets of ZGYCF and 853 POI disease targets, with 231 overlapping targets. Analyzing this network, we found that components such as quercetin, kaempferol, and *β*-sitosterol are closely connected within the network and occupy key positions, indicating that these may be the potential active components of ZGYCF in treating POI. Quercetin is a natural flavonoid compound with moderate estrogenic activity and possesses various pharmacological properties such as antioxidant, anti-inflammatory, antibacterial, and anticancer effects ^[23]^. It also promotes granulosa cell proliferation, the development of primordial follicles into mature follicles, and increases estrogen secretion from granulosa cells, thereby improving ovarian function in rats with premature ovarian failure ^[24, 25]^. Additionally, it can enhance the function of porcine ovarian cells to counteract the damage caused by the environmental pollutant toluene ^[26]^. Kaempferol is a flavonol widely found in various plants, known for its antioxidant properties and its ability to resist apoptosis. According to research by Santos JMS and others, kaempferol can promote the activation of primordial follicles and cell proliferation in vitro ^[27]^. Other studies have shown that kaempferol can increase the synthesis of estradiol and reduce the apoptosis of human ovarian granulosa cells, making it effective in the treatment of POI caused by abnormal apoptosis of granulosa cells ^[28]^. *β*-sitosterol is a common plant sterol that exhibits estrogen-like effects. Research has found that *β*-sitosterol can promote the proliferation of granulosa cells in PCOS and reduce their apoptosis ^[29]^. This indicates that ZGYCF treats POI through a multi-component, multi-target, and multi-pathway approach.

To further explore the mechanism of ZGYCF in the treatment of POI, we constructed a PPI network of common targets between ZGYCF and POI. Through GO and KEGG analyses, we identified the main functions of the related genes and reviewed relevant literature. Tp53, Bcl-2, and Caspase-3 were identified as important targets, with the p53 signaling pathway being a critical pathway. The p53 signaling pathway is mediated by the p53 gene and the p53 protein it encodes. It can detect abnormal signals both inside and outside the cell, such as DNA mutations and damage, and regulate its upstream and downstream targets. By modulating processes like cell cycle, apoptosis, senescence, and DNA repair, the p53 signaling pathway helps to correct cellular metabolic and structural abnormalities^[30]^. The p53 signaling pathway primarily regulates key apoptotic factors such as the caspase family proteins (e.g., caspase-3) and the Bcl-2 protein family (e.g., Bcl-2, Bax) through the intrinsic mitochondrial pathway and the extrinsic death receptor pathway, thereby inducing apoptosis^[31]^. Studies have shown that p53, also known as TP53, can induce apoptosis in ovarian granulosa cells. It is a marker of apoptosis and is located downstream in the apoptosis signaling pathway. Its expression is upregulated when ovarian function declines^[32]^. TP53 is involved in DNA repair and regulates the expression of apoptotic genes^[33]^. When dysfunction occurs in cell cycle regulation during in vitro culture of oocytes, the expression of TP53 and Caspase-3 is upregulated, improving DNA damage repair mechanisms, which in turn leads to cell cycle arrest or apoptosis^[34]^. Additionally, based on the Sankey pathway diagram, three apoptosis-related genes (TP53, Caspase-3, and Bcl-2) are enriched in the p53 signaling pathway. Therefore, regulating apoptosis mediated by the p53 signaling pathway may be a potential mechanism by which ZGYCF treats POI.

To preliminarily verify the active components and molecular mechanisms of ZGYCF in treating POI, we examined quercetin, kaempferol, and *β*-sitosterol against the core targets identified through the PPI network. The results revealed that the binding energies between quercetin, kaempferol, *β*-sitosterol, and the active binding sites of Tp53, Bcl-2, and Caspase-3 were all ≤ -6.5 kJ/mol. This indicates that the core components can spontaneously bind with the target proteins and exhibit strong binding activity, suggesting that Tp53, Bcl-2, and Caspase-3 may be key targets for ZGYCF’s therapeutic effect. To validate the results predicted by network pharmacology and molecular docking, we conducted cell experiments targeting key molecules in the p53 signaling pathway and the molecular docking targets. We selected KGN cells as the research model and used ACR-induced KGN cells to simulate the abnormal granulosa cell apoptosis observed in POI. In our previous studies, we identified 50 μM of ACR as the optimal concentration for intervening in KGN cells^[20]^. In this study, 50 μM ACR was used to treat KGN cells, and the results showed that apoptosis in the ACR group of KGN cells was increased, Bcl-2 expression was decreased, while p53, Bax, Caspase-3 expression, and the Bcl-2/Bax ratio were increased. This indicates that ACR exposure significantly affects human ovarian granulosa cells. After intervention with ZGYCF-containing serum, Bcl-2 expression in KGN cells was increased, while the expression of p53, Bax, Caspase-3, and the Bcl-2/Bax ratio were decreased, with a reduction in granulosa cell apoptosis. These results suggest that ZGYCF-containing serum can reduce abnormal granulosa cell apoptosis induced by ACR, potentially through the regulation of the p53 signaling pathway, which is consistent with the predictions from network pharmacology. This provides scientific evidence supporting the use of ZGYCF in the treatment of POI.

## 6 Conclusion

Through the use of network pharmacology, molecular docking, and cellular experiments, this study has demonstrated that ZGYCF can serve as a reproductive protective agent, capable of preventing, alleviating, and treating ACR-induced apoptosis of human ovarian granulosa cells, which is associated with female reproductive disorders. It enhances the resistance of the female reproductive system to the adverse effects of environmental pollutants, thereby providing therapeutic guidance for women who are occupationally or environmentally exposed to ACR.

## Author Statement

We confirm that neither the manuscript nor any parts of its content are currently under consideration or published in another journal. No conflict of interest exits in the submission of this manuscript. Each of the co-authors has seen and agrees with each of the changes made to this manuscript in the revision. All authors have approved the manuscript and agree with its submission to “*International Immunopharmacology*”.

## CRediT authorship contribution statement

**Xinmiao Zhang:** Writing-original draft, Software, Data curation, Conceptualization. **Hongyan Xi:** Writing-original draft, Software, Data curation, Conceptualization. **Tianyu Gao:** Writing-original draft, Validation, Software. **Shupeng Liu:** Writing-original draft, Visualization, Validation, Software. **Rongxia Li:** Writing-review & editing, Project administration, Funding acquisition, Conceptualization.

## Data availability

Data will be made available on request.

